# Biological network topology is a structural prior for data-efficient sparse learning

**DOI:** 10.64898/2026.03.12.711455

**Authors:** Tousif Jamal, Tansu Celikel

## Abstract

Biological networks, from gene co-expression circuits to cortical connectomes and animal societies, share sparse, modular, small-world topologies thought to reflect optimization for efficient information routing under resource constraints. Whether these structural signatures can act as inductive biases for artificial learning is unknown. We initialized sparse multilayer perceptrons with adjacency matrices inferred by mutual information from molecular, structural, functional, and behavioral networks. We compared them with dense networks and density-matched synthetic graphs on four classification benchmarks. Biologically initialized networks outperformed fully connected baselines, most strongly when training data were scarce: with 100 training examples, a gene co-expression template classified handwritten digits with 65% accuracy, versus 11% for a dense network. Randomly sparsified, rewired, degree-preserved, scale-free, and small-world graphs of matched density did not reproduce this low-data advantage or the low trial-to-trial variance of biological templates. The arrangement of connections, not their number, carries a transferable computational prior.

**HIGHLIGHTS:**

- Sparse networks wired with biological topologies learn from very few examples
- Molecular, brain, and social templates outperform dense baselines
- Density-matched random, scale-free, and small-world graphs do not match them
- Which connections exist, not how many, sets accuracy and stability

**eTOC BLURB:** Jamal and Celikel show that sparse artificial networks wired according to molecular, neural, and social interaction networks learn more accurately and more stably from few examples than dense or density-matched synthetic networks, identifying the arrangement of biological connections, rather than their sparsity, as a transferable computational prior.

**GRAPHICAL ABSTRACT:** 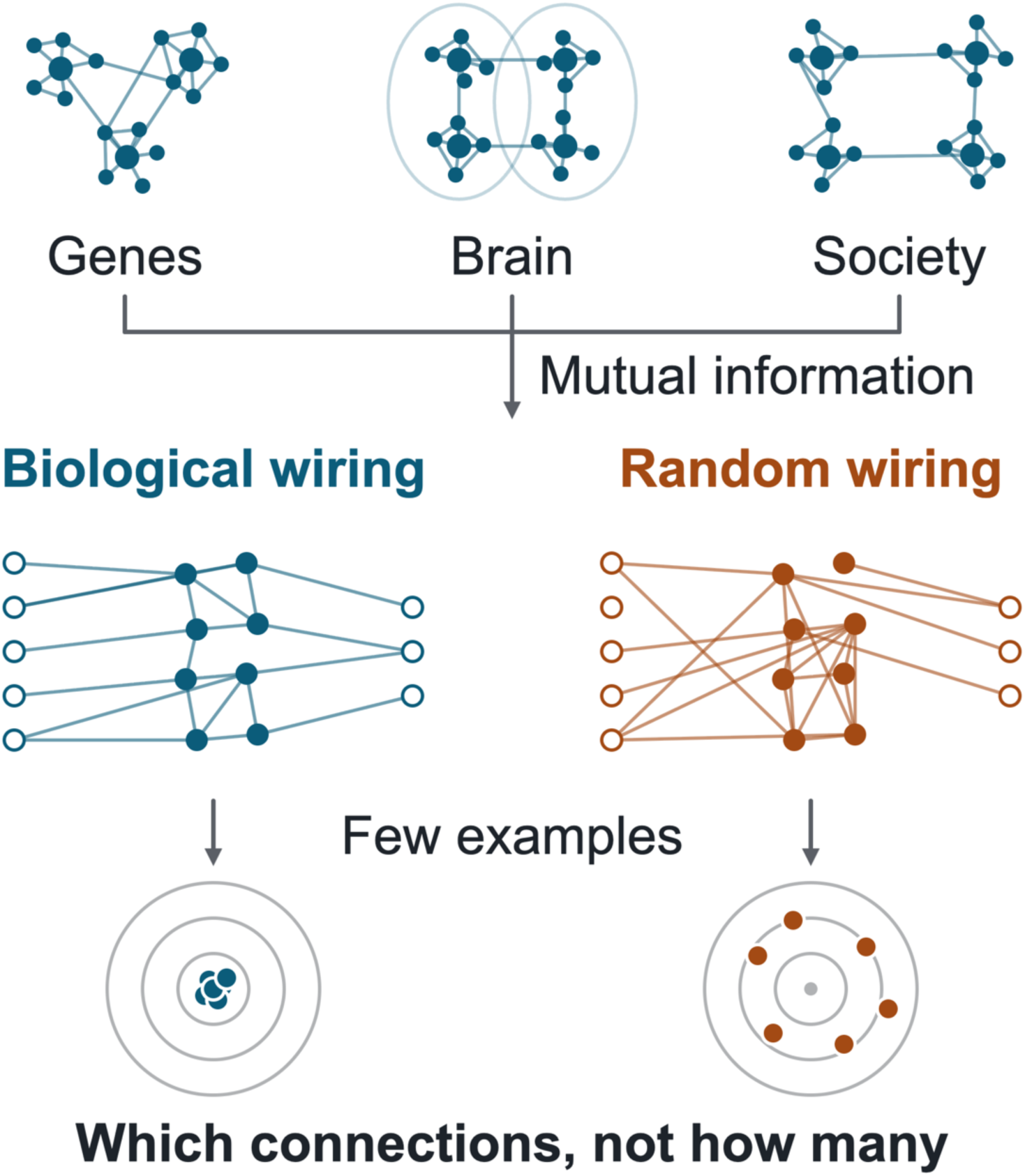

## INTRODUCTION

Animals generalize from remarkably few examples. A young predator learns to detect prey from dozens of encounters; the developing visual cortex calibrates orientation selectivity within a narrow postnatal window; the immune system mounts a specific response on first contact with a novel pathogen. This data efficiency arises not only from biological learning rules but also from the architectures within which learning takes place^1^. Neural circuits, gene regulatory networks, and animal social networks share a recurring structural signature: sparse, modular connectivity with small-world organization, which minimizes wiring cost while supporting information integration^2^. These properties are thought to reflect optimization under two opposing pressures, maximizing adaptive function while minimizing metabolic and developmental cost^2,3^. Within these constraints, biological networks are not fixed. Sensory experience continuously reshapes synaptic, cellular, and molecular organization in a layer-specific and topographically constrained manner^4–10^, so that network structure, and even neuronal identity, is itself shaped by input^11,12^.

Small-world and modular topologies recur across scales and substrates, from cortical connectomes^13,14^ and cellular-resolution circuit maps^15,16^ to gene co-expression^8,10^ and ecological interaction networks^17^. This recurrence suggests a convergent solution to the general problem of routing information efficiently under resource constraints^2,18^. Evolutionary simulations support this view: modular, small-world structure arises spontaneously under selection for performance in variable environments, without explicit architectural constraints^3,19^. Importantly, the structural information in biological networks is not reducible to summary statistics such as connection density or degree distribution. It includes higher-order motifs, community structure, and the placement of hub nodes, each of which may reflect distinct adaptive pressures.

Machine learning likewise relies on inductive biases, structural assumptions built into a model’s architecture, to learn efficiently. Convolutional weight sharing exploits translational invariance, recurrent architectures encode temporal dependencies, and attention mechanisms support flexible relational reasoning. Each of these improves generalization when data are limited. Network science has begun to inform this search. In randomly wired and relational-graph architectures, graph properties such as clustering and path length predict performance^20,21^, and sparse training can evolve scale-free connectivity during learning^22^. Connectome-derived recurrent and reservoir networks have also been tested, with mixed results relative to random wiring^23–25^. Yet most architectural priors are still designed by hand or found by search, leaving the structural solutions produced by biological optimization largely untested^1^.

The lottery ticket hypothesis offers a conceptual bridge. Dense networks contain sparse “winning ticket” subnetworks that, when trained from their original initialization, match the performance of the full network^26^. The specific pattern of initial connectivity, not only its density, therefore determines learning efficiency. Identifying such tickets, however, requires training the dense network first, which negates the efficiency gain. We propose that biological optimization has already solved this search problem: the connection topologies of biological networks are pre-existing, metabolically tested candidate tickets, shaped to generalize from limited experience across the variable environments an organism encounters.

Here we test this hypothesis. We introduce MiPiNet (mutual information pre-initialized networks), a framework that sets the connectivity of sparse multilayer perceptrons (MLPs) from biological adjacency matrices inferred with mutual information^27^. Mutual information captures linear and nonlinear dependencies, and we have previously used it to quantify information transfer in single neurons and sensory circuits^28–30^. We ask whether biological structure, transferred directly to artificial classifiers, acts as a beneficial inductive bias. We compare biologically initialized networks with synthetic alternatives that progressively remove biological structure (randomly rewired, degree-preserved, scale-free, and Watts–Strogatz small-world networks). The comparison spans four benchmarks covering image, categorical, and high-cardinality data, with emphasis on the low-data regime, in which structural priors should matter most.

## RESULTS

We asked whether biologically derived topologies confer learning advantages when used to set the connectivity of sparse MLP classifiers (STAR Methods). Four connectivity matrices served as structural templates: a co-expression network of housekeeping genes (HKG) from mouse somatosensory cortex; structural and functional brain connectivity matrices from the MPI-Leipzig Mind-Brain-Body dataset^31^; and a behavioral association network of bottlenose dolphins (*Tursiops truncatus*)^32,33^. Although these networks span different biological substrates and timescales, all share hallmarks of efficient network organization: sparse connectivity (mean density, approximately 9% of possible edges), small-world organization, and modular structure. All models were evaluated on four benchmarks: Digit Recognition (MNIST)^34^, Objects (Fashion-MNIST)^35^, Selection (Nursery)^36^, and Plants (Plant States)^37^. Training sets ranged from 100 to 10,000 samples, with 50 independent trials per condition. We report mean accuracy ± SD throughout.

We investigated whether biologically derived network topologies confer learning advantages when used to define the connectivity of sparse multilayer perceptron (MLP) classifiers (STAR Methods). Four biologically derived connectivity matrices served as structural templates: a housekeeping gene co-expression network (HKG) from mouse somatosensory cortex; structural and functional brain connectivity networks from the MPI-Leipzig Mind-Brain-Body dataset³⁰,³¹; and a behavioral association network of bottlenose dolphins (*Tursiops truncatus*)³²,³³. These networks represent distinct biological substrates and organizational scales, providing an opportunity to investigate whether biological connectivity patterns influence computational performance across different classification tasks. The structural properties of each network, including node count, edge count, connectivity density, clustering coefficient, characteristic path length, small-world index, and modularity, are summarized in Suplemental file 1.

All architectures were evaluated across four classification benchmarks: Digit Recognition (MNIST)³⁴, Objects (Fashion-MNIST)³⁵, Selection (Nursery)³⁶, and Plants (Plant States)³⁷, using training-set sizes of 100, 300, 1,000, 5,000, and 10,000 samples. Each condition was evaluated across 50 independent experimental trials. Classification performance is reported as mean accuracy ± standard deviation (SD). Statistical differences between architectures were assessed using paired statistical tests across matched experimental trials, with Holm correction for multiple comparisons. In addition to frequentist significance testing, Bayesian factors^38,39^ (BF₁₀) were calculated to quantify the relative evidence supporting the alternative hypothesis over the null hypothesis. Detailed statistical results, including test statistics, corrected p-values, effect sizes, and BF₁₀, are provided in the corresponding Results sections and supplementary file.

### Biological structural priors improve classification, especially when data are scarce

Across four classification benchmarks, biologically initialized networks exhibited network- and dataset-dependent advantages over fully connected architectures, particularly under data-limited conditions (Figure 1). In Digit Recognition, the Behavioral network achieved 67.55% ± 7.43% accuracy with 100 training samples, compared with 20.20% ± 3.97% for its fully connected counterpart (Holm-corrected *p* < 0.001, BF₁₀ = 3.24 × 10³⁵). This advantage persisted at 300 samples (71.57% ± 4.58% versus 46.00% ± 3.78%; *p* < 0.001, BF₁₀ = 2.39 × 10³²). At 100 samples, structural and functional connectivity also significantly improved classification (*p* < 0.001 for both; BF₁₀ = 325.98 and 2.87 × 10³, respectively), while HKG initialization showed a smaller improvement (67.65% ± 8.04% versus 64.55% ± 8.01%; *p* = 0.033, BF₁₀ = 2.37) compared to the others.

**Figure 1.**
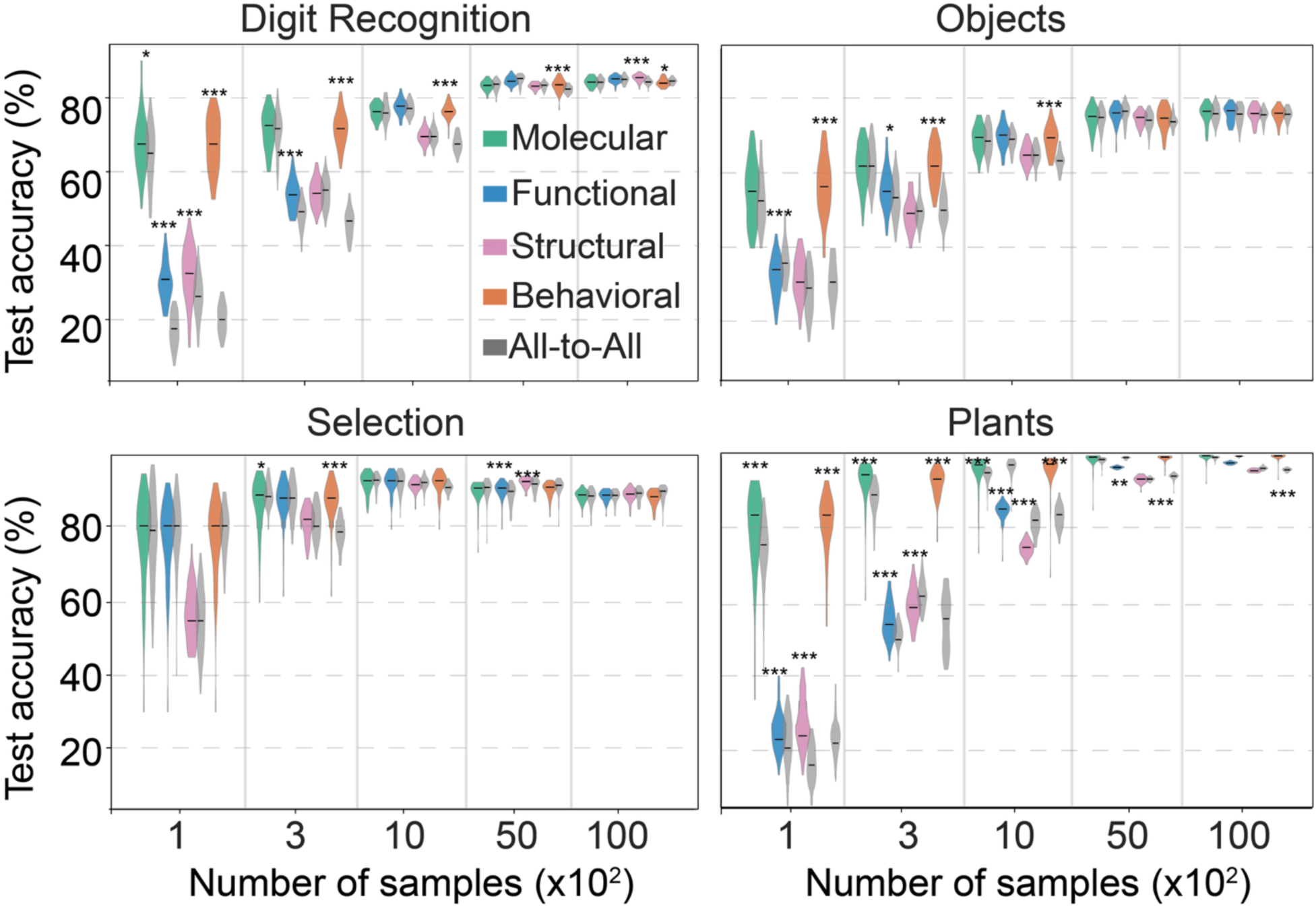
Classification performance of biologically initialized and fully connected networks. Violin plots showing classification accuracy across five training-set sizes (100, 300, 1,000, 5,000, and 10,000 samples) in four classification benchmarks: Digit Recognition, Objects, Selection, and Plants States. Networks initialized with molecular (HKG), structural, functional, and behavioral connectivity templates were compared with their corresponding fully connected (all-to-all) architectures. For each comparison, the fully connected architecture was matched to the corresponding biological network in size (number of nodes), ensuring that differences in classification performance were not attributable to network size. Each violin represents the distribution of classification accuracies across 50 repeated trials per condition. Statistical comparisons were performed using two-sided paired *t*-tests, with Holm correction across the four biological-network comparisons within each dataset and training-set size. Asterisks indicate statistically significant differences (*p* < 0.05, **p** < 0.01, ***p*** < 0.001; Holm-adjusted). Bayesian evidence was quantified using Bayes factors (BF₁₀). Complete statistical results are provided in File S1.

Comparable advantages were observed in Objects and Plants States. With 100 training samples, Behavioral initialization significantly outperformed fully connected architectures in Objects (55.80% ± 7.52% versus 30.40% ± 6.07%; *p* < 0.001, BF₁₀ = 1.02 × 10²¹) and Plants States (80.20% ± 8.51% versus 21.25% ± 6.31%; *p* < 0.001, BF₁₀ = 9.77 × 10³⁴). HKG and structural connectivity also significantly improved classification in Plants States (*p* = 0.003 and *p* < 0.001; BF₁₀ = 11.75 and 5.20 × 10¹⁰, respectively).

In Selection dataset, none of the biological networks significantly outperformed their fully connected counterparts with 100 training samples. However, Behavioral initialization showed a significant advantage at 300 samples (87.50% ± 5.93% versus 78.28% ± 4.06%; *p* < 0.001, BF₁₀ = 6.01 × 10¹¹). Across benchmarks, differences between biological and fully connected architectures generally diminished as training-set size increased, although some network-specific effects persisted (See File S1).

### Sparsity alone does not explain the advantage

To test whether the classification advantages of biological connectivity could be explained solely by reduced network density, we systematically sparsified fully connected architectures across nine sparsity levels (10%–95%) and compared their performance with biologically initialized networks (Figure 2 and Figure S1). Repeated-measures ANOVAs revealed significant effects of sparsification across most datasets and training-set sizes, although the magnitude of these effects depended on the network architecture. For the fully connected counterparts of the Behavioral network, sparsification significantly affected classification with 100 training samples in Digit Recognition (*F*(9, 441) = 209.67, partial η² = 0.811), Objects (*F*(9, 441) = 130.46, partial η² = 0.727), Selection (*F*(9, 441) = 47.69, partial η² = 0.493), and Plants States (*F*(9, 441) = 154.85, partial η² = 0.760; all *p* < 0.001).

**Figure 2.**
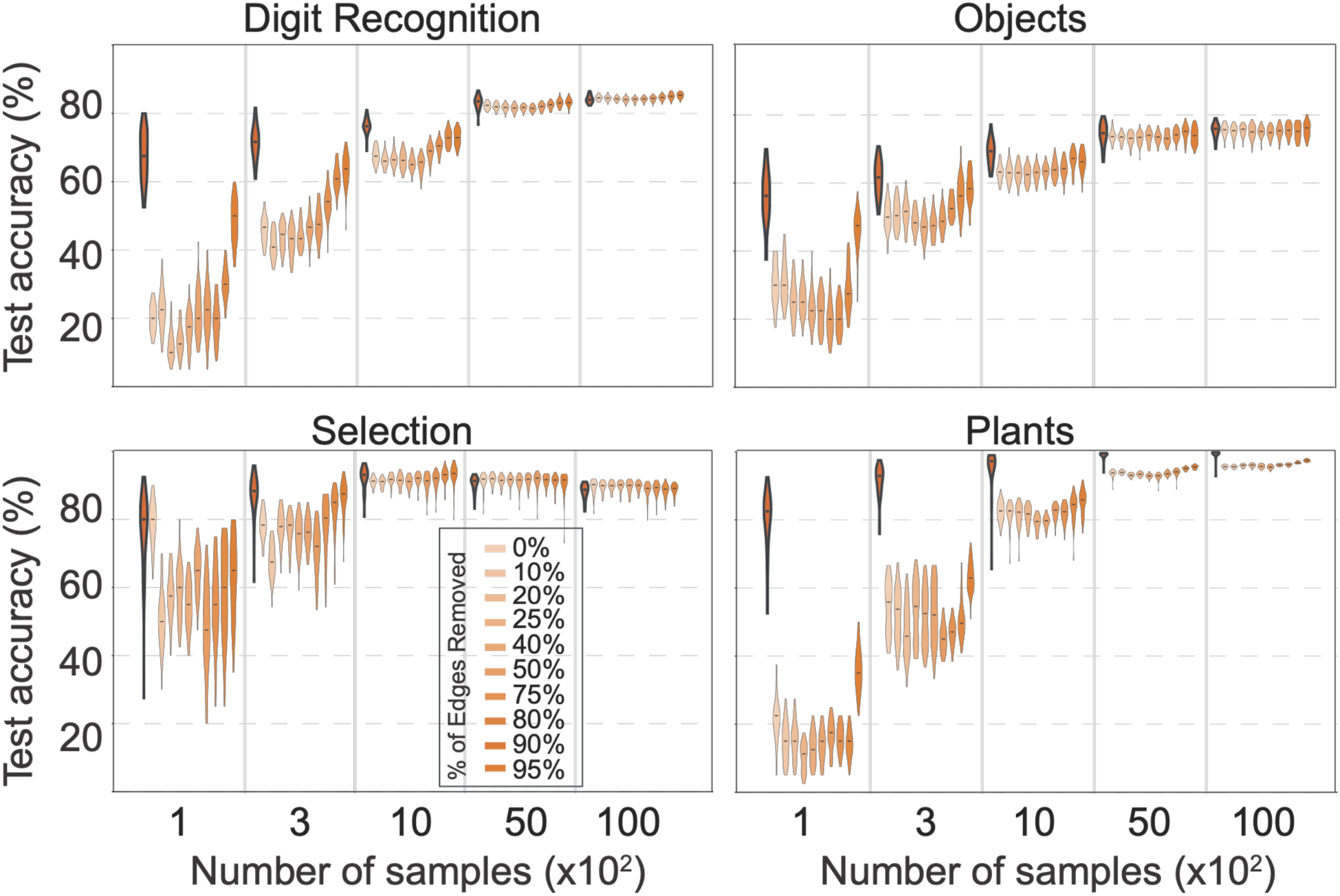
Sparsity alone does not explain the classification advantages of biological network topologies. Violin plots showing classification accuracy of randomly sparsified fully connected networks (0%–95% edge removal) compared with the corresponding biological behavioral network (highlighted in bold). All fully connected and sparsified variants were matched to the behavioral network in size (181 nodes), ensuring that differences in classification performance were not attributable to network size. Sparsification levels are represented by varying color opacity. Performance was evaluated across four classification benchmarks (Digit Recognition, Objects, Selection, and Plants States) and five training-set sizes (100, 300, 1,000, 5,000, and 10,000 samples). Each violin represents the distribution of classification accuracies across 50 independent trials per condition. Vertical lines separate training-set sizes. Corresponding analyses for the molecular, structural, and functional connectivity networks are presented in Figure S1.

Despite these effects, Behavioral connectivity significantly outperformed highly sparsified fully connected networks under data-limited conditions. With 100 training samples, Behavioral initialization achieved higher accuracy than the 95%-sparsified controls in Digit Recognition (67.55% ± 7.43% versus 48.35% ± 6.44%; Holm-corrected *p* < 0.001, BF₁₀ = 1.24 × 10¹⁶), Objects (55.80% ± 7.52% versus 45.90% ± 6.01%; *p* < 0.001, BF₁₀ = 6.55 × 10⁵), and Plants States (80.20% ± 8.51% versus 35.75% ± 6.29%; *p* < 0.001, BF₁₀ = 4.07 × 10³¹). A similar advantage was observed in Selection, where Behavioral connectivity also outperformed the strongest sparsified control (77.10% ± 11.06% versus 65.00% ± 6.70%).

The effects of sparsification differed considerably across the remaining biological networks (Figure S1). With 100 training samples, sparsification had relatively limited effects on HKG-associated fully connected architectures, with significant differences observed only in Selection (*F*(9, 441) = 2.84, partial η² = 0.055, *p* = 0.0058) and Plants States (*F*(9, 441) = 16.06, partial η² = 0.247, *p* < 0.001). In contrast, structural and functional network controls showed significant effects of sparsification across all four datasets (structural: *F*(9, 441) = 3.23–1093.00, partial η² = 0.062–0.957; functional: *F*(9, 441) = 33.93–1229.00, partial η² = 0.409–0.962; all Greenhouse–Geisser-corrected *p* < 0.05). When biological architectures were included in the comparisons, significant differences persisted for structural and functional connectivity across all four datasets, whereas HKG showed significant differences only in Selection and Plants States. Across training-set sizes, these heterogeneous responses indicate that the effects of sparsification depend strongly on the underlying network architecture and dataset.

Sparsity is therefore necessary but not sufficient: which connections are present, not merely how many, determines the benefit. Random edge removal discards the structure encoded in biological connectivity, and having few connections is not equivalent to having the right ones.

### Biological topology outperforms progressively matched synthetic graphs

To check whether the classification advantages of biological connectivity depended on specific network organization, we compared the original biological architectures with randomly rewired, degree-preserved shuffled, and hub-connected alternatives, as well as small-world networks with varying rewiring probabilities (Figures 3, S2 and S3). Repeated-measures ANOVAs revealed significant effects of alternative connectivity structures on classification accuracy, with substantial differences across biological networks and datasets. These architectural manipulations also produced distinct patterns of trial-to-trial variability.

**Figure 3.**
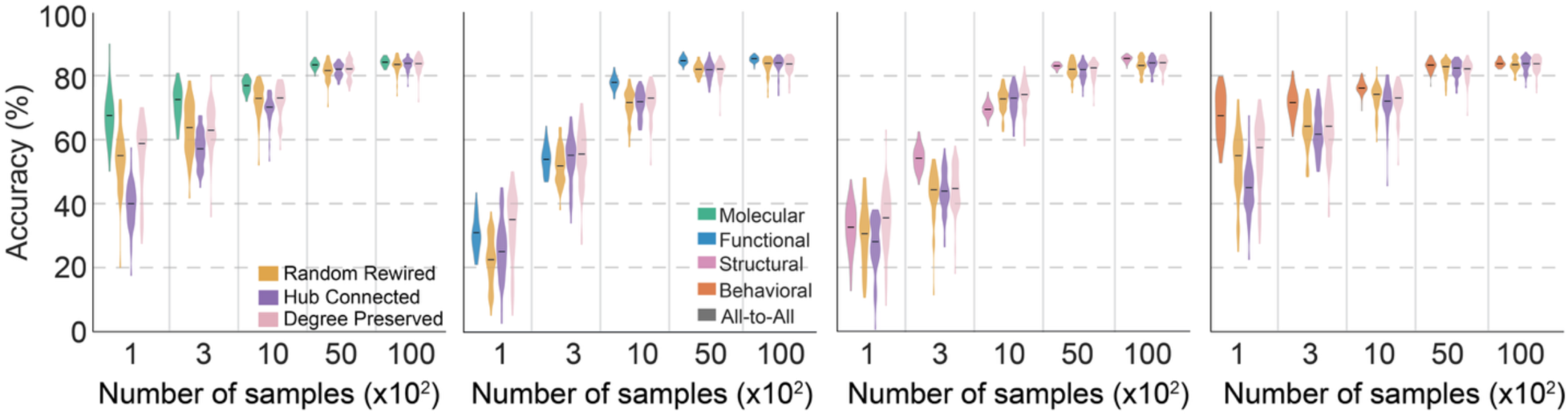
Network organization influences classification performance beyond connectivity density. Violin plots showing classification accuracy of networks initialized with four biological connectivity templates: Molecular (HKG), Functional, Structural, and Behavioral, compared with their corresponding randomly rewired, hub-connected (Barabási–Albert), and degree-preserved shuffled counterparts. Performance was evaluated on Digit Recognition (MNIST) across five training-set sizes (100, 300, 1,000, 5,000, and 10,000 samples). Each violin represents the distribution of classification accuracies across 50 repeated trials per condition, with colors distinguishing network architectures. Vertical lines separate training-set sizes. See Figure S2 for corresponding comparisons across the other three classification benchmarks (Objects, Selection, and Plants States).

Behavioral connectivity exhibited particularly strong architectural sensitivity. With 100 training samples, classification performance differed significantly among the original, randomly rewired, degree-preserved shuffled, and hub-connected architectures in Digit Recognition (*F*(3, 147) = 250.73, partial η² = 0.837), Objects (*F*(3, 147) = 42.74, partial η² = 0.466), Selection (*F*(3, 147) = 5.23, partial η² = 0.096), and Plants States (*F*(3, 147) = 580.22, partial η² = 0.922; all Green-house–Geisser-corrected *p* < 0.05). In Digit Recognition, Behavioral connectivity achieved 67.55% ± 7.43% accuracy, significantly outperforming the hub-connected architecture (40.50% ± 6.29%; Holm-corrected *p* < 0.001, BF₁₀ = 3.11 × 10²³), while performing comparably to random rewiring (64.75% ± 8.05%; *p* = 0.080, BF₁₀ = 1.63). In Plants States, Behavioral connectivity also significantly outperformed hub-connected (36.90% ± 6.48%; *p* < 0.001, BF₁₀ = 1.67 × 10²⁸) and randomly rewired networks (76.60% ± 6.99%; *p* = 0.006, BF₁₀ = 14.85). Although Behavioral connectivity did not consistently exhibit the lowest SD, its classification advantages were maintained despite comparable variability across repeated trials.

Small-world connectivity similarly influenced classification performance in Digit Recognition (*F*(4, 196) = 146.26, partial η² = 0.749), Objects (*F*(4, 196) = 45.76, partial η² = 0.483), and Plants States (*F*(4, 196) = 202.49, partial η² = 0.805; all Greenhouse–Geisser-corrected *p* < 0.001), but not Selection (*p* = 0.094). In Digit Recognition, Behavioral connectivity significantly outperformed all four small-world configurations (all Holm-corrected *p* < 0.001; BF₁₀ = 6.43 × 10¹⁵–1.89 × 10²⁵), while exhibiting a similar range of variability (SD = 7.43% versus 7.36%–7.76%). Comparable classification advantages were observed in Objects and Plants States, where Behavioral connectivity achieved higher mean accuracy without a corresponding reduction in variability. In Selection, small-world architectures achieved comparable classification performance, with some configurations exhibiting lower SD than the original biological network.

The remaining biological networks showed heterogeneous responses (Figure S2). Alternative connectivity structures significantly affected structural and functional network performance across all four datasets with 100 training samples (structural: *F*(3, 147) = 37.50–63.36, partial η² = 0.434–0.564; functional: *F*(3, 147) = 106.86–1154.48, partial η² = 0.686–0.959; all Green-house–Geisser-corrected *p* < 0.001). Comparable effects were observed for small-world connectivity (structural: *F*(4, 196) = 42.72–191.32, partial η² = 0.466–0.796; functional: *F*(4, 196) = 61.02–630.00, partial η² = 0.555–0.928; all *p* < 0.001). In contrast, HKG connectivity showed no significant overall differences across alternative or small-world architectures at 100 training samples (all *p* > 0.05). Variability likewise depended on the biological source and network transformation, indicating that lower trial-to-trial variation was not a universal property of biological connectivity.

Together, these results indicate that classification performance depends on network organization beyond connectivity density alone. Behavioral connectivity showed particularly strong advantages over hub-connected and small-world alternatives under data-limited conditions, generally without increased variability relative to these controls.

### Biological structural priors persist when networks are scaled up

Biological networks are inherently constrained by the size of their source datasets. To determine whether their classification advantages extend beyond these original dimensions, we systematically expanded all the biological network to 2×, 3×, 4×, and 8× its original size using a degree-preserving algorithm. This approach stochastically introduced additional nodes while preserving network sparsity, degree distribution, and weight statistics (STAR Methods; Figure 4 and Supplementary Figures). Across all four classification benchmarks, biologically informed initialization generally maintained classification performance under moderate upscaling (2×–4×), whereas performance at 8× was more variable. These findings suggest that the benefits of biological structural priors are not confined to the dimensions of the original network and can extend to larger architectures, although their effectiveness may depend on network size and the availability of training data.

**Figure 4.**
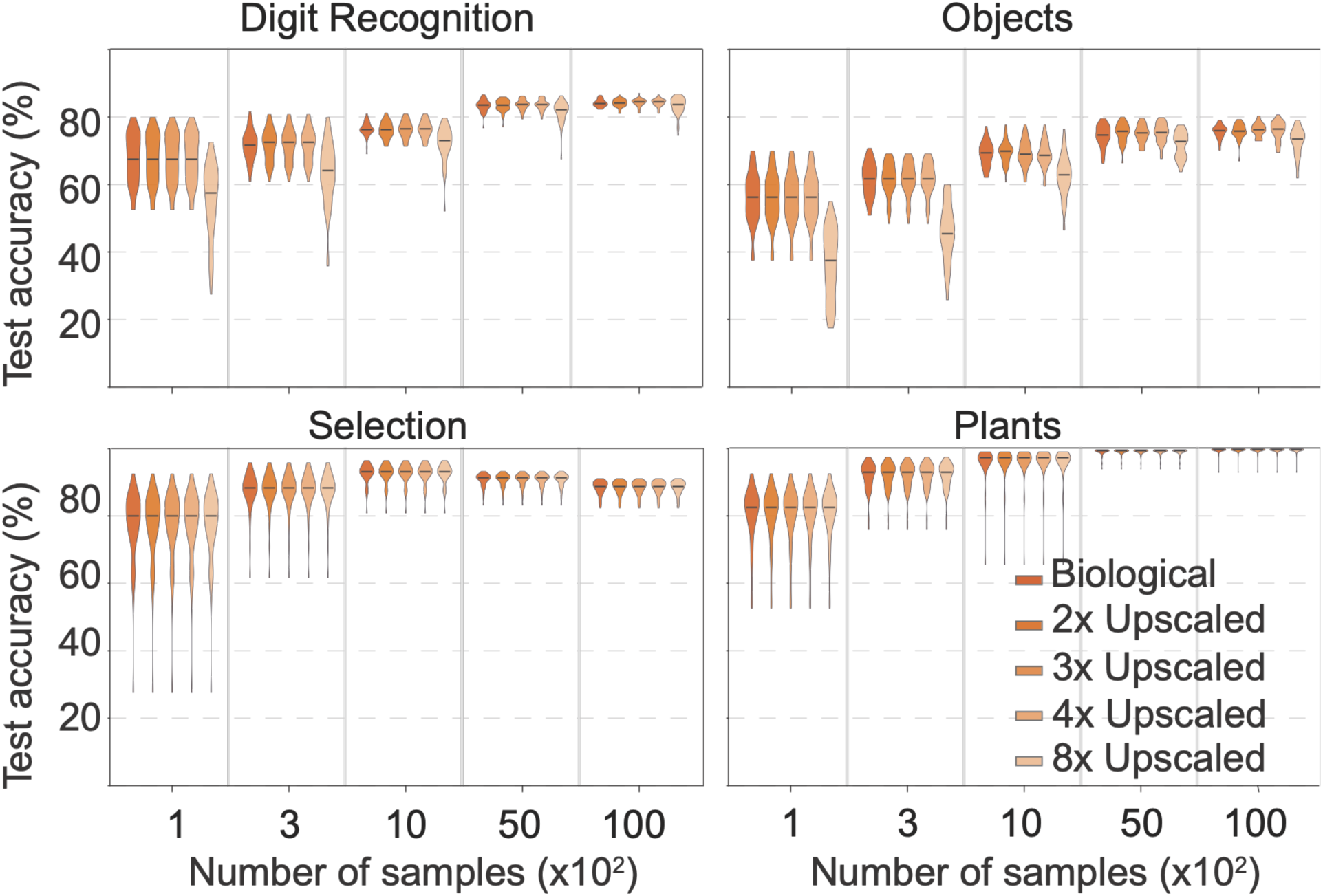
Expanded biological networks retain classification performance across increasing network sizes. Classification accuracy of the original Behavioral network and its expanded counterparts (2×, 3×, 4×, and 8× the original network size) across four classification benchmarks: Digit Recognition, Objects, Selection, and Plants States. Networks were expanded using a degree-preserving algorithm that stochastically introduces additional nodes while maintaining network sparsity, degree distribution, and weight statistics (STAR Methods). Performance was evaluated across five training-set sizes (100, 300, 1,000, 5,000, and 10,000 samples). Data represent mean ± SD across 50 repeated trials per condition. Corresponding analyses for the molecular, structural, and functional connectivity networks are presented in Figure S4.

This stability was particularly evident for Behavioral connectivity, which maintained identical classification accuracy across the original, 2×, 3×, and 4× architectures in all four datasets. With 100 training samples, accuracy remained at 67.55% ± 7.43% in Digit Recognition, 55.80% ± 7.52% in Objects, 77.10% ± 11.06% in Selection, and 80.20% ± 8.51% in Plants States. In Selection and Plants States, performance remained unchanged even at 8× scaling, with no variation across scaling conditions. Although repeated-measures ANOVAs revealed significant overall effects in Digit Recognition (*F*(4, 196) = 57.52, partial η² = 0.540) and Objects (*F*(4, 196) = 120.47, partial η² = 0.711; both Greenhouse–Geisser-corrected *p* < 0.001), these effects were attributable to differences at the highest scaling factor rather than changes across the moderate scaling conditions.

Comparable stability was observed across the remaining biological networks (Figure S3). HKG, structural, and functional connectivity maintained unchanged classification performance across all scaling conditions in Selection and Plants States. Accordingly, no ANOVA could be computed for these datasets because the trial-level accuracies were identical across scales. In Objects, structural connectivity also showed no significant effect of upscaling (*F*(4, 196) = 1.74, partial η² = 0.034, *p* = 0.193). Although significant overall effects were detected for HKG in Digit Recognition (*F*(4, 196) = 46.67, partial η² = 0.488) and Objects (*F*(4, 196) = 116.43, partial η² = 0.704), structural connectivity in Digit Recognition (*F*(4, 196) = 178.64, partial η² = 0.785), and functional connectivity in Digit Recognition (*F*(4, 196) = 401.39, partial η² = 0.891) and Objects (*F*(4, 196) = 33.65, partial η² = 0.407; all *p* < 0.001), these differences primarily reflected changes at the largest scaling factor.

Importantly, 8× upscaling represents a substantial increase in network size relative to the original architectures. Under limited training-data conditions, such expansion may increase the demands on network optimization and data availability, potentially contributing to the variable performance observed at this scale. Nevertheless, the preservation of classification accuracy across moderate scaling conditions, together with the absence of detectable scaling effects in several benchmarks, indicates that biological connectivity-based initialization can maintain its performance characteristics as network dimensions increase. These results suggest that biological structural priors are compatible with network expansion, although their effectiveness at substantially larger scales may depend on the availability of sufficient training data.

## DISCUSSION

Biological network topology acts as a beneficial inductive bias for sparse artificial classifiers. Networks initialized with molecular, structural, functional, and behavioral adjacency matrices outperformed fully connected baselines, randomly sparsified networks, and synthetic structural approximations on all benchmarks and in most data conditions. The largest gains came when training data were scarce. That templates from such different substrates conferred similar advantages is consistent with the hypothesis that biological optimization converges on structural solutions to the problem of learning efficiently from limited data.

### The active ingredient is organization, not sparsity

Randomly rewired networks share the sparsity and weight distribution of their biological counterparts, and degree-preserved networks share their degree sequence. Neither reproduced biological performance in the low-data regime, so connection density alone cannot explain the advantage. This dissociation parallels evolutionary simulations in which modular and small-world topologies arise under selection for performance in variable environments, even from random starting points^3,19^. What distinguishes biological from randomly sparse networks is not a generic property of sparse graphs but a specific arrangement that supports efficient information routing. Higher-order features, including local clustering, modular community structure, and hub-and-spoke connectivity, provide inductive biases that sparsity or degree distribution alone cannot recover. Biological networks must generalize from limited experience^2^, and our results suggest that this property can be transferred to artificial systems as an architectural prior. Connectome-instantiated recurrent and reservoir networks have performed on par with, but not better than, randomly wired networks^24,25^. The feedforward, low-data setting studied here, in which structural priors should matter most, may explain this difference.

### Convergent organization and the limits of structural approximation

Watts–Strogatz networks reproduce the mean topological statistics of biological networks but not their specific arrangement. Their convergence toward biological performance with large training sets confirms that clustering and path length carry genuine structural information, but convergence was incomplete in the low-data regime. Biological networks likely contain information absent from generative small-world models. This includes the identity of highly connected hubs, the configuration of community boundaries, and heterogeneous weights shaped by Hebbian-like developmental and experience-dependent processes^4,6,42^. This residual advantage matters most when labeled data are scarce, mirroring the conditions under which biological networks must operate.

The recurrence of small-world and modular topology across substrates suggests that these properties reflect general principles of efficient computation under resource constraints rather than task-specific adaptations^2,13,14^. Our results add that the specific realization of these principles matters: the history of a biological network encodes structure that generic models do not fully replicate. Libraries of biological networks are therefore an underexplored source of structural priors. These include connectomes from different species and brain regions, regulatory networks from different cell types, and interaction networks from different habitats. Conversely, the performance of a sparse learner initialized with a given topology offers a quantitative assay of that topology’s computational value. Such an assay could compare biological networks across substrates, species, and experiential histories.

### Biological networks as pre-selected lottery tickets

The lottery ticket hypothesis posits that dense networks contain sparse winning subnetworks that can be found by magnitude-based pruning after full training^26^. MiPiNet can be viewed as supplying candidate tickets at the outset, bypassing dense training. Gradient descent searches for sparse subnetworks within a single training run. Biological optimization, by contrast, has searched the space of topologies over many generations and across far more variable challenges than any benchmark. The result is a set of sparse connectivity patterns that generalize across computational contexts, as reflected in the task-general advantage reported here. The strong performance of biologically initialized networks in the low-data regime supports this view, because pruning-based methods are least effective there: the dense model itself trains poorly. Combining biological priors with learned rewiring of sparse connectivity^22^ is a natural extension.

### Design principles for neuromorphic and edge computing

More broadly, biological networks are an underused library of structural priors for artificial architectures. The scaling results show that this library is not limited to the size of the source network: templates can be expanded while preserving key organizational properties, with performance recovering as training data increase. Machine learning increasingly targets resource-constrained settings such as edge devices, mobile inference, neuromorphic hardware^41^, and scientific domains with scarce labels. In these settings, biologically grounded initialization offers a training-efficient alternative to post hoc pruning or costly architecture search. Sparse, biologically structured connectivity is also a natural fit for event-driven, spiking implementations of sensory pathways^42,43^. The consistency of the advantage across tasks, and across genomic, cortical, and social templates, suggests that the underlying principles are robust.

The choice of template may also be tunable to the task, with different biological substrates offering different inductive biases. Priors derived from individual neuroimaging or transcriptomic data are a particularly promising direction, combining population-level organization with individual connectivity patterns.

### Limitations of the study

Several limitations should be noted. First, we tested only fixed-dimensional classification; whether biological structural priors transfer to sequential, structured, or multimodal prediction remains to be established. Second, network expansion relied on degree-preserving randomization, which disrupted local motifs and degraded low-data performance. Expansion algorithms that preserve higher-order features, such as triadic closure and community structure, may avoid this cost. Third, although biological topology provides a beneficial inductive bias, ablations that isolate the contribution of specific features (clustering, path length, hub connectivity, or community structure) are needed for a mechanistic account. Fourth, the templates are statistical dependency networks inferred from population-level data and therefore average over individual variability. Our design cannot distinguish whether their structure reflects evolutionary optimization, developmental processes, or experience-dependent refinement. Finally, we compared biological templates with dense and synthetic sparse baselines, not with dynamic sparse training or pruning methods.

## STAR★METHODS

## KEY RESOURCES TABLE

| REAGENT or RESOURCE | SOURCE | IDENTIFIER |
|---|---|---|
| **Deposited data** |  |  |
| MPI-Leipzig Mind-Brain-Body MRI data | Babayan et al.<sup>31</sup> | https://doi.org/10.1038/sdata.2018.308; OpenNeuro: [ac-cession] |
| Mouse L2/3 single-cell RNA-seq (housekeeping genes) | https://barrelomics.sci-ence.ru.nl/ | https://barrelomics.sci-ence.ru.nl/transcriptome |
| Bottlenose dolphin association net-work | Gazda et al.<sup>33</sup>; ASNR<sup>32</sup> | https://bansal-lab.github.io/asnr/ |
| MNIST | LeCun et al.<sup>33</sup> | http://yann.lecun.com/exdb/mnist/ |
| Fashion-MNIST | Xiao et al.<sup>34</sup> | https://github.com/zalan-doresearch/fashion-mnist |
| Nursery | Rajkovic<sup>35</sup> | https://doi.org/10.24432/C5P88W |
| Plants | Hämäläinen<sup>36</sup> | https://doi.org/10.24432/C5HS40 |
| **Software and algorithms** |  |  |
| MiPiNet | This paper | Zenodo: [DOI] |
| NETSCOPE | Bergmans et al.<sup>27</sup> | https://doi.org/10.1101/2022.04.18.488630 |
| NetworkX | Hagberg et al.<sup>45</sup> | https://networkx.org |
| Python | Python Software Founda-tion | https://www.python.org |
| [Deep learning framework, e.g., PyTorch] | https://pytorch.org/ | https://pytorch.org/ |

## RESOURCE AVAILABILITY

### Lead contact

Requests for further information and resources should be directed to and will be fulfilled by the lead contact, Tansu Celikel.

### Materials availability

This study did not generate new materials.

### Data and code availability

- This paper analyzes existing, publicly available data. Accession numbers and DOIs for these datasets are listed in the key resources table.
- All original code, including the MiPiNet framework and the scripts used to generate the synthetic networks and figures, has been deposited at Zenodo and is publicly available as of the date of publication at [DOI].
- Any additional information required to reanalyze the data reported in this paper is available from the lead contact upon request.

## EXPERIMENTAL MODEL AND STUDY PARTICIPANT DETAILS

### Human participants

We collected previously published MRI data from healthy adults aged 20–30 years in the MPI-Leipzig Mind-Brain-Body database^30^ Data collection was approved by the ethics committee of the medical faculty of the University of Leipzig, and all participants gave written informed consent, as described in the original publication^30^.

### Mice

Single-cell transcriptomes were obtained from layer 2/3 of the somatosensory cortex of male and female mice by our group and made publicly available previously^46,47^. No animals were handled for this study.

### Bottlenose dolphins

The behavioral network was derived from published observational data on a free-ranging population of bottlenose dolphins (*Tursiops truncatus*) in Cedar Key, Florida^32^; no animals were handled for this study.

## METHOD DETAILS

### Biological network templates

#### Structural and functional brain connectivity

Structural and functional brain connectivity networks were obtained from the MPI-Leipzig Mind-Brain-Body dataset^31^ comprising 318 unique participants across two partially overlapping neuroimaging protocols: Leipzig Study for Mind–Body–Emotion Interactions (LEMON) and Neuroanatomy & Connectivity (N&C). The LEMON protocol included 228 participants, whereas the N&C protocol included 199 participants, with 109 individuals participating in both. To reduce computational demands, we selected 136 participants sequentially according to participant identifiers (sub-010001 to sub-010136). We used the preprocessed neuroimaging data provided with the dataset, for which the original preprocessing procedures, including relevant artifact correction and confound removal, are described in the accompanying publications. Functional connectivity was estimated from the preprocessed resting-state fMRI time series using the NETSCOPE mutual-information toolbox^27^, whereas structural connectivity was obtained from the diffusion MRI-derived connectivity data. The resulting connectivity matrices were represented as undirected networks, comprising 183 nodes for functional connectivity and 181 nodes for structural connectivity. These networks were subsequently used as biologically derived structural priors in the classification experiments.

#### Housekeeping gene network

Housekeeping genes (HKGs) are expressed in most cell types, support essential cellular functions, and show relatively stable expression across tissues, developmental stages, and experimental conditions, which is why they are widely used as reference genes.

To construct the HKG network, we used single-cell RNA sequencing data from layer 2/3 (L2/3) of the mouse somatosensory cortex, including samples from male and female mice^8^. The dataset comprised transcript counts for 451 genes. After identifying genes shared between male and female samples, we used NETSCOPE^27^ to estimate pairwise mutual information between genes and reconstruct a network based on their statistical dependencies. Weak connections were excluded to retain the strongest gene–gene associations, and disconnected nodes (genes without any remaining connections) were subsequently removed. Consequently, the final network contained 401 nodes compared to the original 451 measured genes. The resulting network retained approximately 9% of all possible connections.

#### Behavioral interaction network

The behavioral connectivity template was derived from the travelling-association network of bottlenose dolphins (*Tursiops truncatus*) in Cedar Key, Florida³³. The network was obtained from the Network Data Repository, which incorporates data from the Animal Social Network Repository³². The original network comprised 188 individuals and represented an undirected, weighted network of associations during travelling behavior, recorded through 5-minute survey scans over 124 sampling days. Edge weights quantified the strength of pairwise associations using the half-weight index. The travelling network was selected from the four activity-specific networks and the overall association network reported in the original study. Following NETSCOPE analysis and the removal of nodes with no nonzero-weight connections, the resulting network comprised 181 nodes and was used as the biological connectivity template for classification.

### Synthetic comparison networks

#### Randomly rewired networks

To separate topology from sparsity and weight distribution, we shuffled the weights of each biological matrix and reassigned them to source–target pairs sampled uniformly across the matrix. The resulting matrix **A′** satisfies ǁ**A′**ǁ₀ = ǁ**A**ǁ₀, preserving sparsity and the marginal weight distribution while removing topological structure. This network serves as the null model for biological structure (Figure S3).

#### Hub-connected (scale-free) networks

Scale-free networks were generated with the Barabási–Albert preferential attachment model^40^. Each new node connects to *m* existing nodes with probability proportional to their degree, producing a heavy-tailed degree distribution with emergent hubs. The value of *m* was chosen to match the total edge count of the biological reference (Figure S3).

#### Degree-preserved shuffled networks

Network connections were randomized using undirected double-edge swaps or directed edge swaps where supported, with a target of min(5E, 20,000) swaps, where E represents the number of edges. Successful swaps preserved the original binary degree distribution while altering network connectivity. However, the original assignment of edge weights to specific connections was not preserved. (Figure S5).

#### Small-world networks

To investigate the contribution of small-world topology to classification performance, synthetic networks were generated using the Watts–Strogatz model implemented in NetworkX⁴⁵ (watts_strogatz_graph(n, k, β)). For each biological network, a regular ring lattice was constructed with the same number of nodes (n) and an approximately matched mean degree (k ≈ 2m/n, rounded to an even integer). Each node was initially connected to its k nearest neighbors, after which edges were randomly rewired with probability β ∈ {0.01, 0.1, 0.3, 0.6}. Rewiring preserved the total number of edges while progressively introducing long-range connections and reducing lattice regularity, without allowing self-connections or duplicate edges. Edge weights were independently sampled from the empirical distribution of nonzero weights in the corresponding biological network, and adjacency matrices were symmetrized to maintain undirected connectivity. These four variants allowed us to assess the effects of network organization while approximately matching network size, connectivity, and weight distribution (Figure S3).

#### Network scaling

To investigate whether biologically informed initialization remains effective at larger network sizes, we expanded the original HKG network by factors of 2, 3, 4, and 8 using a degree-preserving replication and rewiring algorithm (Figure S3). All networks were represented as undirected, weighted adjacency matrices.

First, the original N × N adjacency matrix was replicated along the diagonal of a larger matrix to generate s identical, initially disconnected network copies, where s represents the scaling factor. The resulting expanded network contained sN nodes and retained the original connectivity patterns and edge weights within each copy.

Second, connections between adjacent network copies were introduced through degree-preserving double-edge swaps. For each pair of adjacent copies, two existing edges, one from each copy, were randomly selected. These edges were removed and replaced with two cross-copy connections, provided that the proposed connections did not already exist and did not introduce self-loops. The original edge weights were transferred to the newly established connections.

Third, this rewiring procedure was repeated until approximately 3% of the original network’s edges had been exchanged between each pair of adjacent copies, with a minimum of one successful swap per pair. The number of attempted swaps was limited to the greater of 200 or 100 times the target number of successful swaps.

This procedure preserved the degree of every node, the total number of edges, and the distribution of nonzero edge weights while introducing connections between replicated network modules. The expanded adjacency matrices were subsequently used to initialize classifiers trained and evaluated using the same procedures as the original networks (Figure S3).

#### Fully connected baselines

Fully connected (all-to-all) networks were initialized with dense weight matrices (W ∈ ℝᴺˣᴷ), where N represents the input dimension and K the number of output classes. Initial weights were independently sampled from a standard normal distribution, N(0, 1). Randomly sparsified controls used the same initialization procedure, with a proportion of connections initially set to zero. To match the input dimensionality to the network size, datasets with fewer than N features were zero-padded, whereas datasets with more than N features underwent principal component analysis (PCA). The number of retained principal components was determined by the network size, subject to the available sample and feature dimensions. Any remaining dimensions were zero-padded. All architectures were evaluated using identical training procedures and hyperparameters (Figure S5).

### MiPiNet architecture and training

To evaluate the effects of biological network organization on classification performance, we used a single-layer multiclass perceptron initialized with biological or synthetic connectivity matrices. For each network, the original N × N adjacency matrix was mapped onto an input-to-output weight matrix by selecting the columns corresponding to the output classes. Input features were aligned with the resulting input dimension using principal component analysis (PCA) when dimensionality reduction was required, or zero-padding when the number of features was smaller than the input dimension.

For biologically initialized networks, the numerical adjacency values, including mutual-information-derived weights where applicable, were used directly as initial connection weights. Zero-valued entries were initially retained as zero, but no fixed binary connectivity mask was imposed during training. Thus, both initially present and absent connections could be modified by the learning algorithm. Fully connected and synthetic network controls were evaluated using the same classification architecture and training procedure.

The classifier consisted of a single linear input-to-output layer without hidden layers or nonlinear activation functions. Class predictions were obtained by selecting the output with the highest linear score. Training used a mistake-driven perceptron learning rule: for each incorrectly classified sample, the weights associated with the correct class were increased by the input vector multiplied by the learning rate, while those associated with the incorrectly predicted class were decreased by the same amount. No weight update was performed for correctly classified samples.

All architectures were trained using identical hyperparameters: a learning rate of 0.01, a batch size of one sample, and a single training epoch. No separate loss-based optimizer, regularization procedure, or early-stopping criterion was applied. Data were divided into training (60%) and test (40%) sets, and classification performance was evaluated across 50 repeated trials using different random seeds. The same learning procedure was applied to all biological and synthetic network architectures to enable direct comparison of their classification performance.

### Benchmark datasets

#### Digit Recognition

MNIST^34^ comprises 70,000 grayscale images of handwritten digits (28 × 28 pixels; 60,000 training and 10,000 test images; 10 classes), flattened to 784-dimensional input vectors.

#### Objects

Fashion-MNIST^35^ comprises 70,000 grayscale images of clothing (28 × 28 pixels; 60,000 training and 10,000 test images; 10 classes).

#### Selection

The Nursery dataset^36^ contains 12,960 instances described by eight categorical features (parental occupation, childcare arrangements, family structure, financial standing, and social and health conditions). It was derived from a hierarchical decision model for ranking applications to nursery schools in Ljubljana, Slovenia.

#### Plants

The Plant States dataset^37^ contains 34,781 records describing the geographic distribution of plant species and genera across U.S. states and Canadian provinces. Each record contains a Latin plant name and the geographic regions in which it occurs. We converted the transactional data into a binary feature matrix using 70 geographic indicators, where each feature denotes the presence or absence of a plant in a particular state or province. For classification, the 20 most frequently occurring plant names were assigned individual class labels, while all remaining names were grouped into an additional “Other” category, yielding 21 output classes. The resulting task evaluated whether geographic occurrence patterns could predict plant identity.

## QUANTIFICATION AND STATISTICAL ANALYSIS

Each network architecture was evaluated across five training-set sizes (100, 300, 1,000, 5,000, and 10,000 samples), with 50 repeated trials per condition. Trials used different random seeds to generate training and test splits, while identical seeds were used across architectures to ensure paired comparisons. Data were divided into training (70%) and held-out test (30%) sets, and classification accuracy was calculated as the proportion of correctly classified test observations. Results are reported as mean ± SD across trials. Differences between architectures were assessed using two-sided paired *t*-tests with Holm correction for multiple comparisons within predefined comparison families. Effect sizes were quantified using Cohen’s dz, and Bayesian evidence was evaluated using Bayes factors (BF₁₀) with a Jeffreys–Zellner–Siow Cauchy prior (r = 1/√2). Statistical significance was defined as Holm-adjusted *p* < 0.05. In all plots, small-world network performance is displayed with transparency proportional to the rewiring probability (β).

To assess differences in classification performance across network architectures, we performed one-way repeated-measures analyses of variance (RM-ANOVAs), treating network architecture as a within-trial factor and trial identity as the repeated-measures unit. Analyses were conducted separately for each benchmark dataset, biological network source, and training-set size. Only trials with complete observations across all compared architectures were included. Greenhouse–Geisser corrections were applied to account for potential violations of sphericity, and effect sizes were quantified using partial eta squared (η²ₚ).

Pairwise comparisons between biological, fully connected, and alternative network architectures were performed using two-sided paired *t*-tests, matching observations by trial identity. Statistical significance was assessed at α = 0.05, with *p*-values adjusted for multiple comparisons using the Holm step-down procedure within each predefined comparison family, dataset, biological network source, and training-set size. Effect sizes were reported as Cohen’s *d* with subscript z (dz), calculated as the mean paired difference divided by its standard deviation. Mean differences and their 95% confidence intervals were calculated from the paired observations.

Bayesian evidence for pairwise differences was assessed using two-sided Bayesian paired *t*-tests with a Jeffreys–Zellner–Siow Cauchy prior (scale parameter *r* = 1/√2). Bayes factors (BF₁₀) quantified the relative evidence supporting the alternative hypothesis (H₁) over the null hypothesis (H₀), with values greater than 1 favoring a difference between conditions and values below 1 favoring the absence of a difference. Following Kass and Raftery^38,39^, BF₁₀ values of 1–3, 3–20, 20–150, and greater than 150 were interpreted as weak, positive, strong, and very strong evidence for H₁, respectively.

Bayesian analyses were complementary to frequentist hypothesis testing and were not subjected to Holm correction. All statistical analyses were performed in Python using SciPy and custom analysis scripts. Complete descriptive statistics, paired comparisons, effect sizes, Holm-adjusted *p*-values, and Bayes factors are provided in the Supplementary Tables.

## ACKNOWLEDGMENTS

This work was funded by the European Union’s Horizon 2020 research and innovation program under Marie Skłodowska-Curie (grant agreement no. 860949), the European Regional Development Fund (MIND, no. 122035), and the Netherlands Organisation for Scientific Research

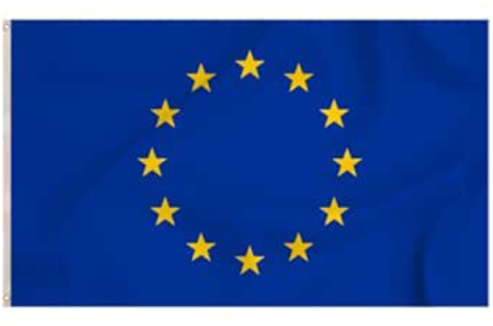

(NWO-ALW Open Competition, no. 824.14.022).

## AUTHOR CONTRIBUTIONS

Conceptualization, T.J. and T.C.; methodology, T.J. and T.C.; software, T.J.; formal analysis, T.J.; investigation, T.J.; writing - original draft, T.J. and T.C.; writing - review & editing, T.J. and T.C.; supervision, T.C.; funding acquisition, T.C.

## DECLARATION OF GENERATIVE AI AND AI-ASSISTED TECHNOLOGIES

During the preparation of this work, the authors used Claude (Anthropic) to edit the manuscript for clarity and journal formatting. After using this tool, the authors reviewed and edited the content as needed and take full responsibility for the content of the publication.

## SUPPLEMENTAL INFORMATION

Supplemental information can be found online at [DOI].

---

**Table S1.**
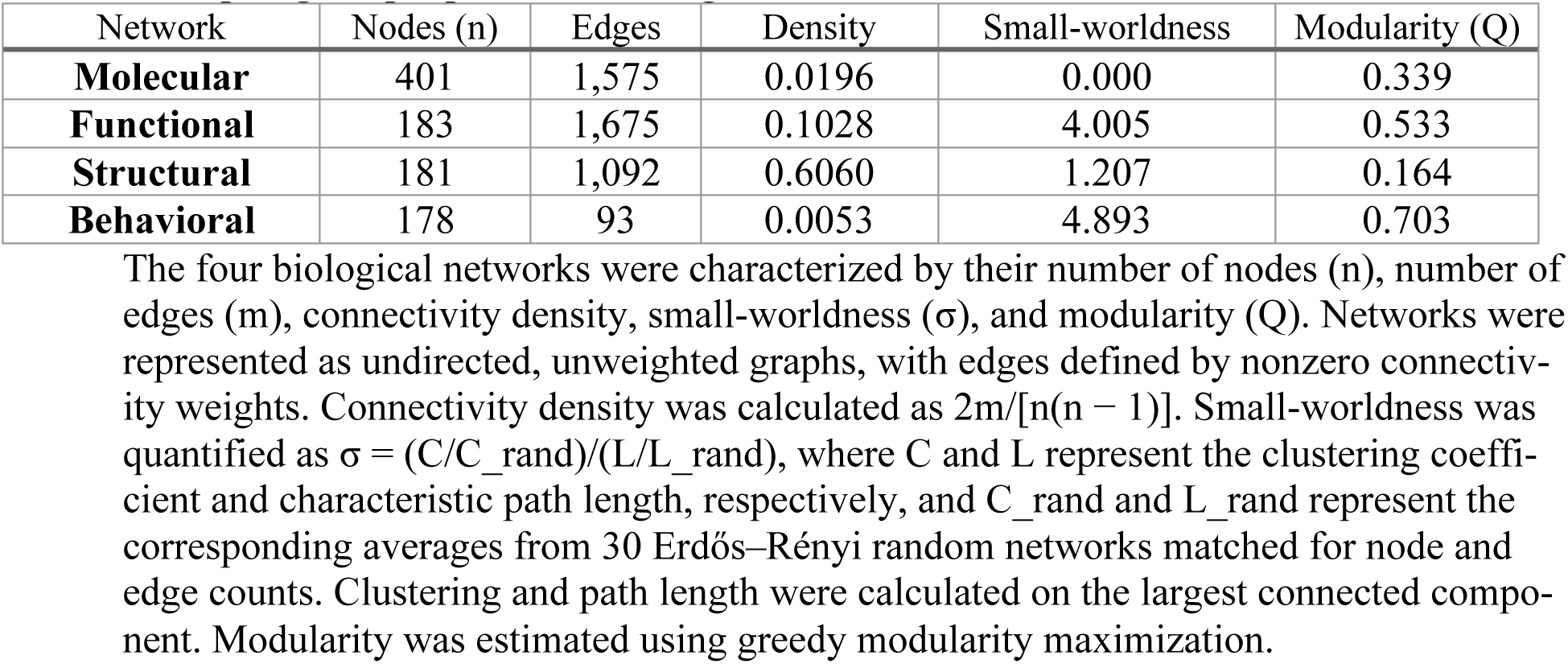
Topological properties of biological networks used in MiPiNet.

| Network | Nodes (n) | Edges | Density | Small-worldness | Modularity (Q) |
|---|---|---|---|---|---|
| **Molecular** | 401 | 1,575 | 0.0196 | 0.000 | 0.339 |
| **Functional** | 183 | 1,675 | 0.1028 | 4.005 | 0.533 |
| **Structural** | 181 | 1,092 | 0.6060 | 1.207 | 0.164 |
| **Behavioral** | 178 | 93 | 0.0053 | 4.893 | 0.703 |

The four biological networks were characterized by their number of nodes (n), number of edges (m), connectivity density, small-worldness (σ), and modularity (Q). Networks were represented as undirected, unweighted graphs, with edges defined by nonzero connectiv-ity weights. Connectivity density was calculated as 2m/[n(n − 1)]. Small-worldness was quantified as σ = (C/C\_rand)/(L/L\_rand), where C and L represent the clustering coeffi-cient and characteristic path length, respectively, and C\_rand and L\_rand represent the corresponding averages from 30 Erdős–Rényi random networks matched for node and edge counts. Clustering and path length were calculated on the largest connected compo-nent. Modularity was estimated using greedy modularity maximization.

**Figure S1.**
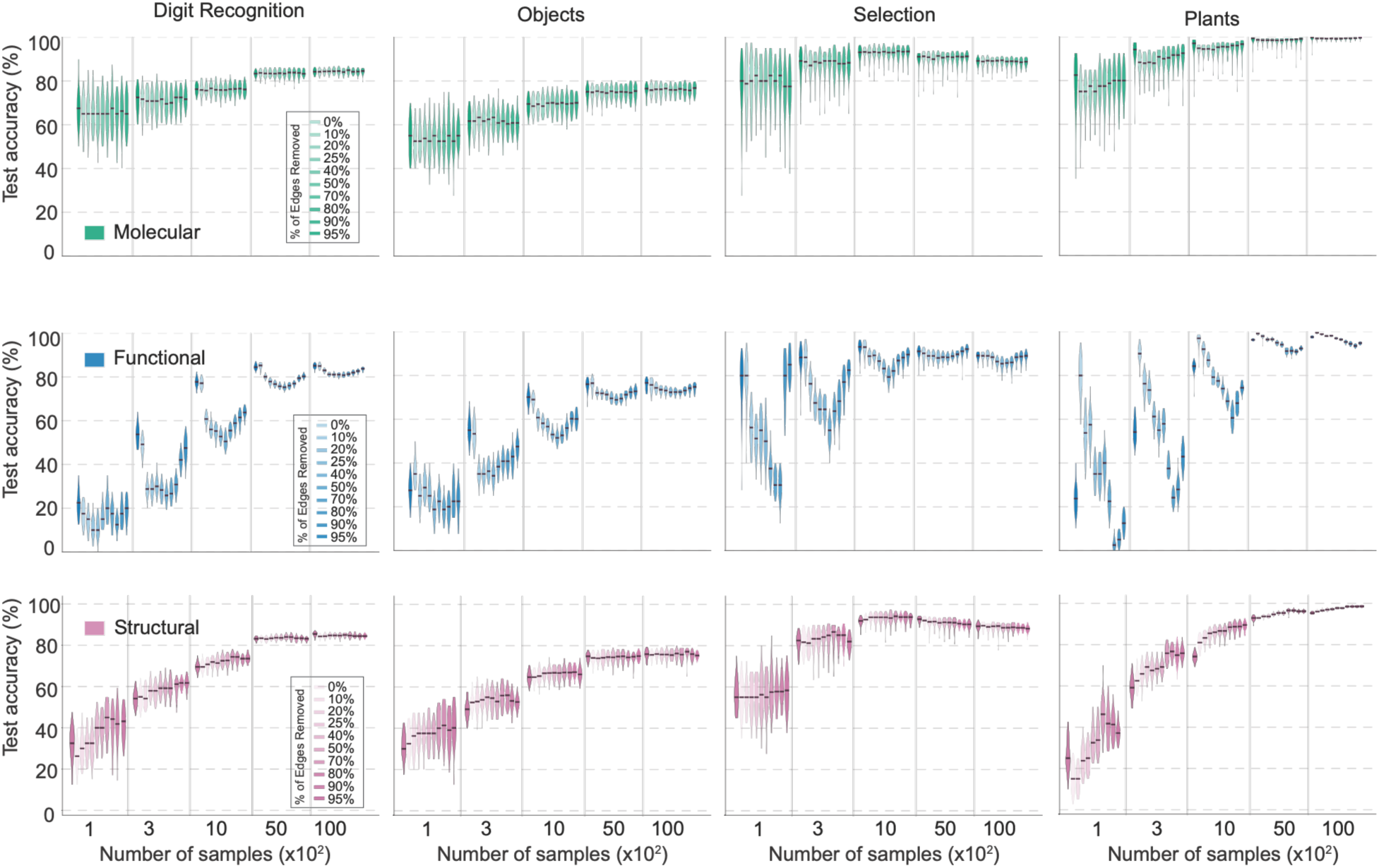
Effects of random sparsification on fully connected networks, related to Figure 2. Violin plots showing classification accuracy of randomly sparsified fully connected networks (0%–95% edge removal) compared with their corresponding biological connectivity templates: molecular (HKG), structural, and functional networks (highlighted in bold). For each comparison, all fully connected and sparsified variants were matched to the corresponding biological network in size (number of nodes), ensuring that differences in classification performance were not attributable to network size. Sparsification levels are represented by varying color opacity. Performance was evaluated across four classification benchmarks (Digit Recognition, Objects, Selection, and Plants States) and five training-set sizes (100, 300, 1,000, 5,000, and 10,000 samples). Each violin represents the distribution of classification accuracies across 50 independent trials per condition. Vertical lines separate training-set sizes. The corresponding analysis for the behavioral network is presented in Figure 2.

**Figure S2.**
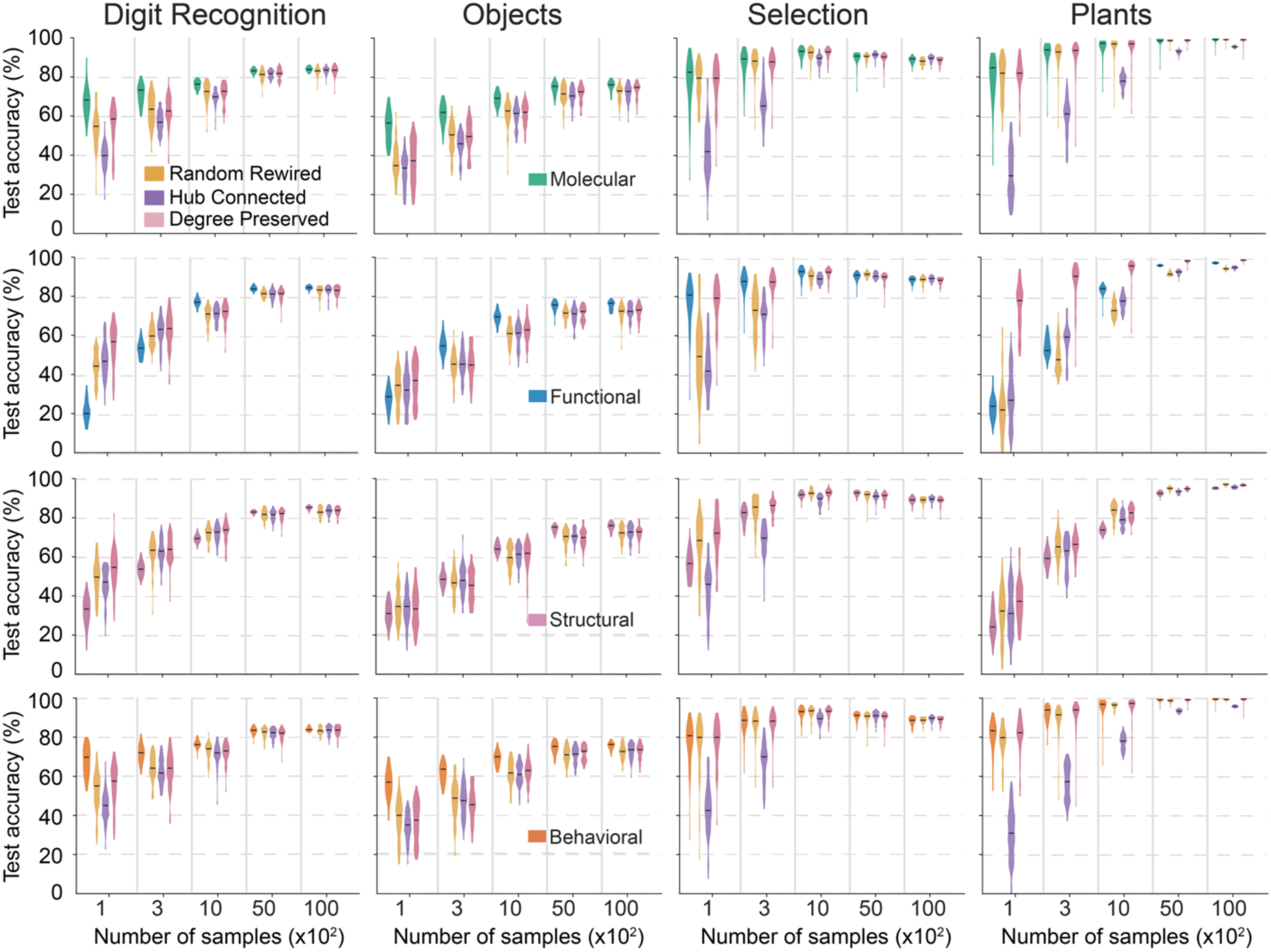
Biological topologies outperform density-matched synthetic graphs across tasks, related to Figure 3. Classification accuracy as a function of training-set size on all four benchmarks for fully connected networks, the four biological templates, and their randomly rewired, hub-connected, and degree-preserved counterparts. Data are mean ± SD across 50 independent trials per condition.

**Figure S3.**
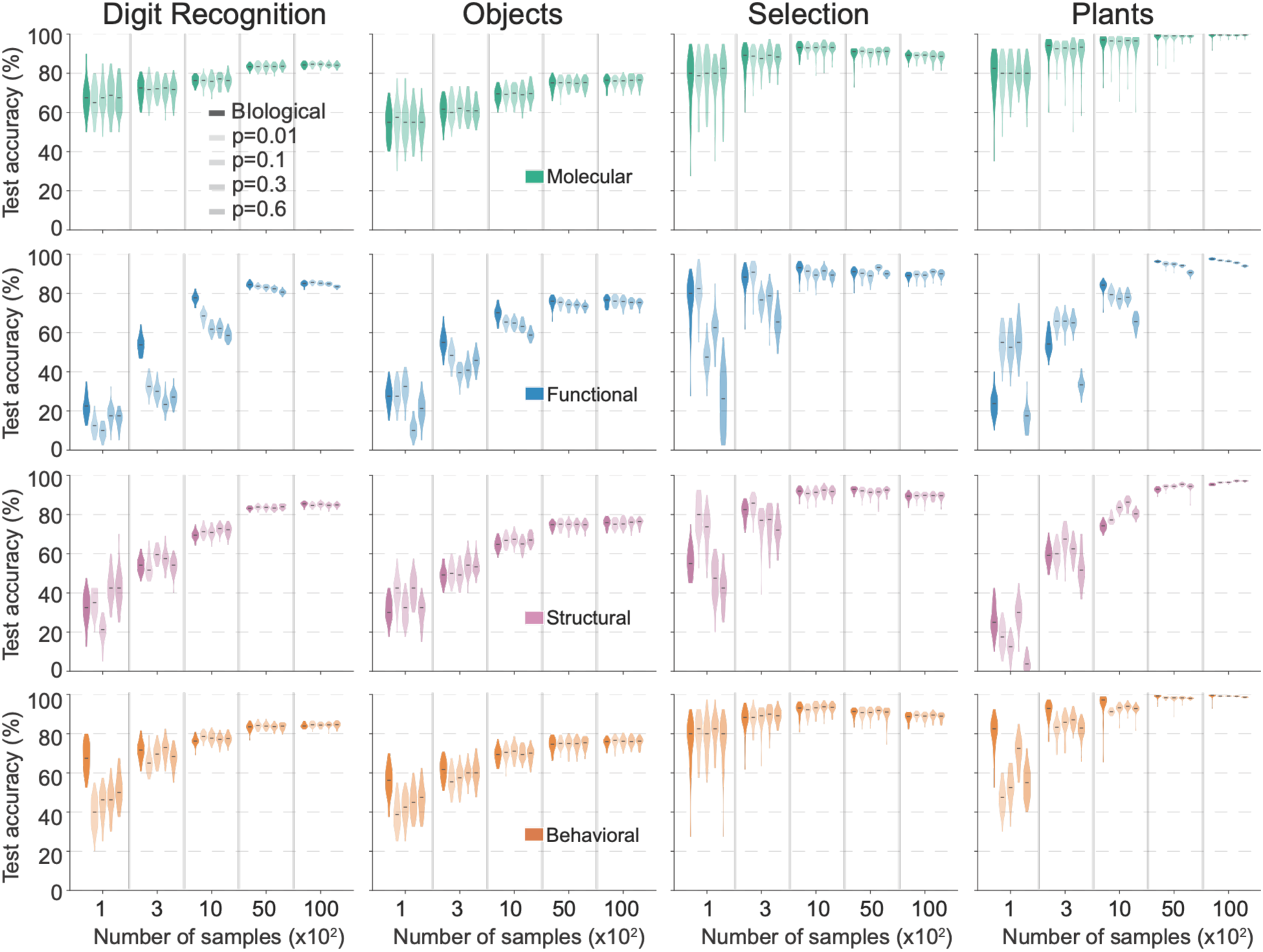
Biological topologies compared with Watts–Strogatz small-world networks, related to Figure 2. Classification accuracy of the biological templates and of Watts–Strogatz networks with rewiring probability β = 0.01, 0.1, 0.3, and 0.6 (transparency scales with β). Data are mean ± SD across 50 independent trials per condition.

**Figure S4.**
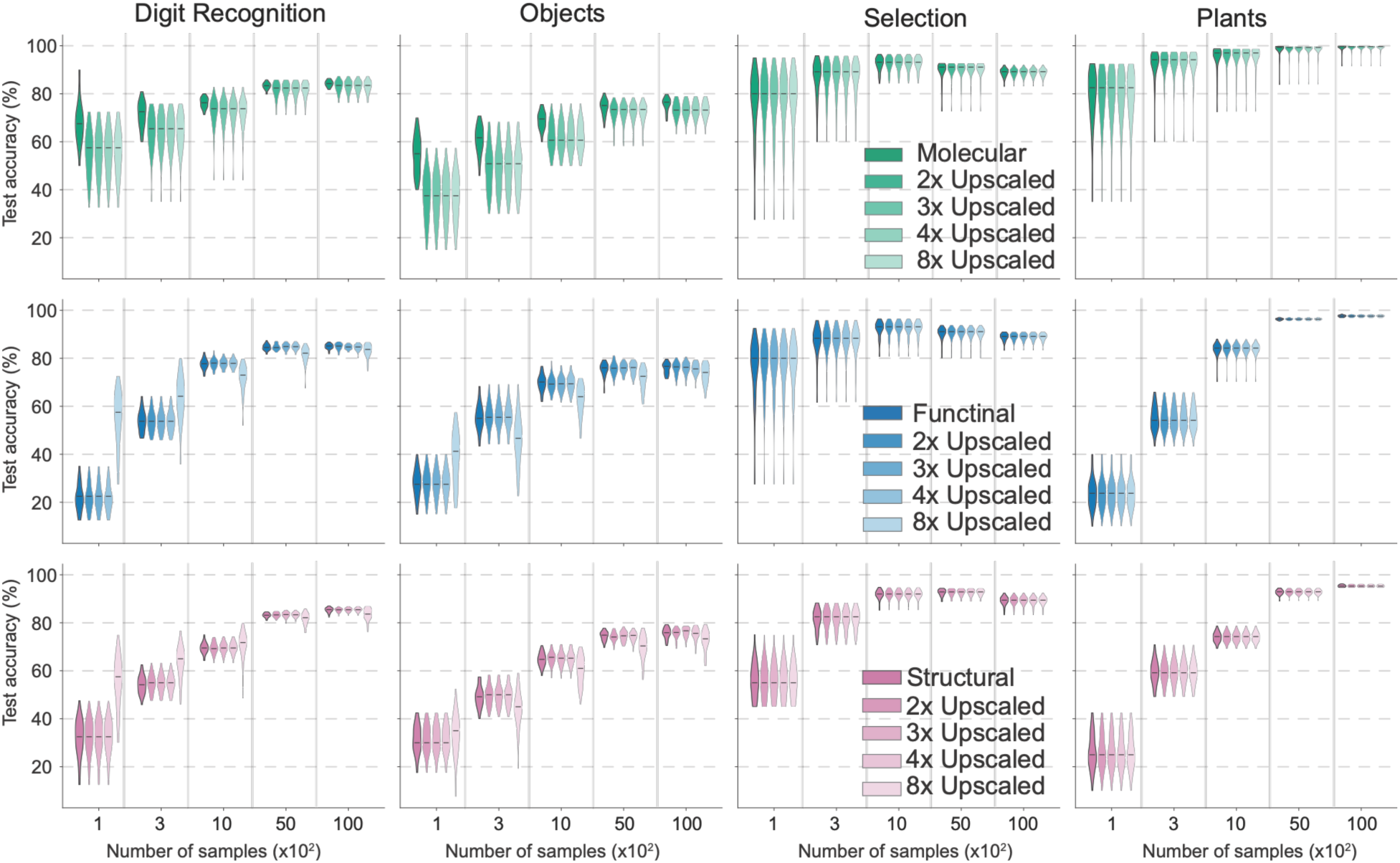
Effects of network expansion on molecular, structural, and functional connectivity networks, related to Figure 4. Classification accuracy of the original molecular (HKG), structural, and functional connectivity networks and their expanded counterparts (2×, 3×, 4×, and 8× the original network size) across four classification benchmarks: Digit Recognition, Objects, Selection, and Plants States. Networks were expanded using the same degree-preserving algorithm as in Figure 4, maintaining network sparsity, degree distribution, and weight statistics (STAR Methods). Performance was evaluated across five training-set sizes (100, 300, 1,000, 5,000, and 10,000 samples). Data represent mean ± SD across 50 repeated trials per condition. The corresponding analysis for the behavioral network is presented in Figure 4.

**Figure S5.**
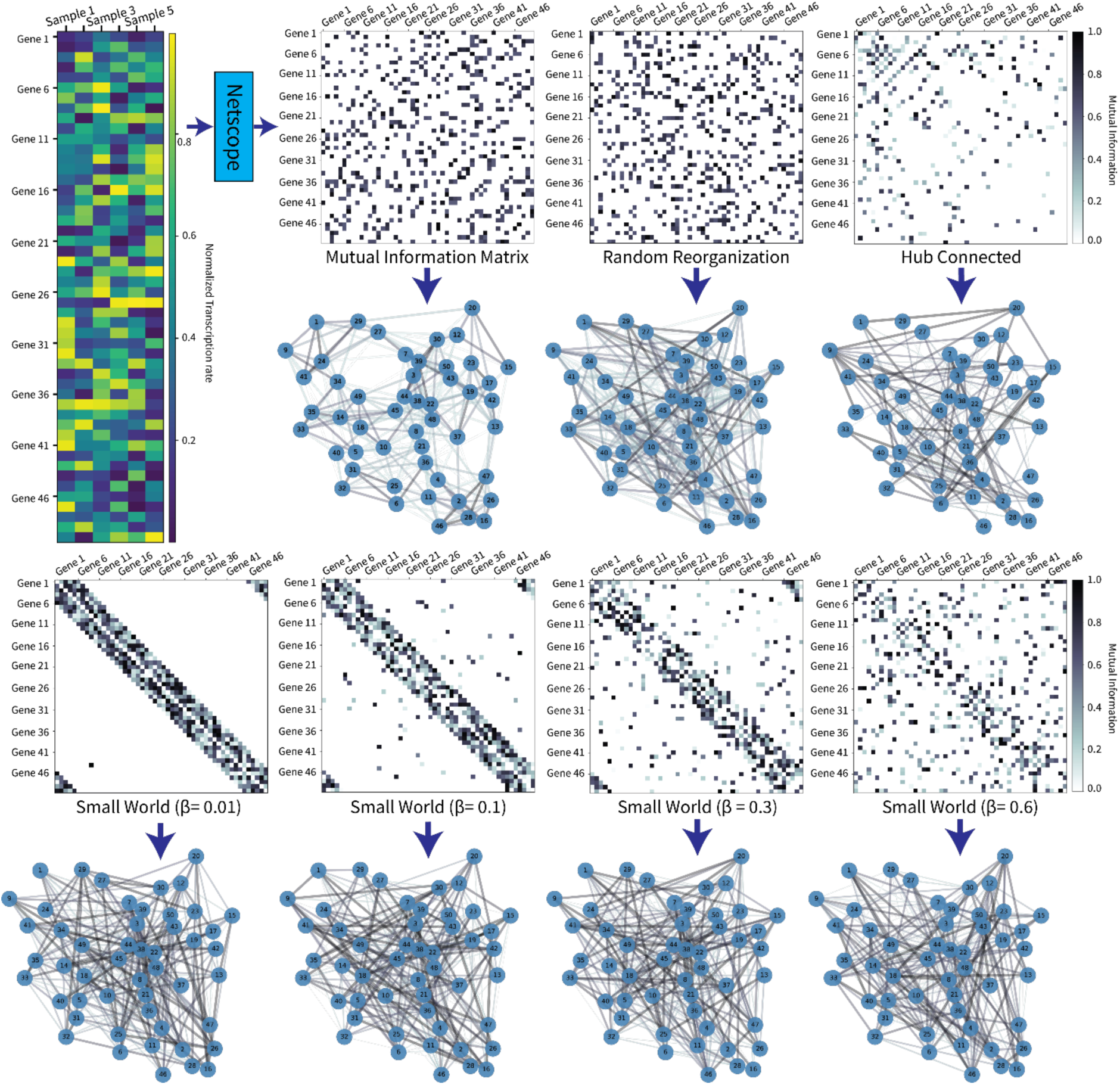
Construction of biological and synthetic network templates, related to Figures 1 and 2. Starting from the gene transcript count matrix (top left), NETSCOPE^26^ computes pairwise mutual information between genes to construct a sparse co-expression network, which serves as the baseline connectivity. Subsequent panels show the synthetic variants used to test the role of topology: (i) random reorganization, which preserves weight distribution and sparsity but randomizes edge placement; (ii) hub-connected (Barabási–Albert) networks with preferential attachment and emergent hubs; (iii) degree-preserved shuffling, which retains node degrees while randomizing connections; and (iv) Watts–Strogatz small-world networks with rewiring probability p, which trade off clustering and path length.

